# Directly selecting differentially expressed genes for single-cell clustering analyses

**DOI:** 10.1101/2023.07.26.550670

**Authors:** Zihao Chen, Changhu Wang, Siyuan Huang, Yang Shi, Ruibin Xi

**Author notes:** These authors contributed equally to this work (Zihao Chen, Changhu Wang).

## Abstract

In single-cell RNA sequencing (scRNA-seq) studies, cell-types and their associated marker genes are often identified by clustering and differential expression gene (DEG) analysis. scRNA-seq data contain many genes not relevant to cell-types and gene selection procedures are needed for more accurate clustering. An ideal gene selection procedure should select all DEGs between cell-types for best cell-type identification. However, because cell-types are unknown, gene selection and DEG analysis are performed separately using different methods. Genes are selected using surrogate criteria not directly related with clustering, which often miss important genes or select unimportant genes. Clustering accuracy could be seriously influenced because of the inferior gene selection. DEGs are often detected by comparing different clusters, leading to many false DEGs due to the selection bias problem. In this paper, we present Festem, a unified method for gene selection and DEG analysis in scRNA-seq studies. Festem investigates gene’s clustering information based on the observation that marginal distributions of DEGs are mixtures of their different cell-type-conditional distributions, and can directly select the clustering-informative DEGs and avoid the selection bias problem. Extensive simulation and real data analyses show that Festem achieves high precision and recall for DEG detection, and enables more accurate clustering and cell-type identification. Applications to several scRNA-seq datasets demonstrate that Festem can identify cell-types that are often missed by other methods. In a large intrahepatic cholangiocarcinoma dataset, we identify CD8+ T cell-types and find that their marker genes are novel prognostic biomarkers.

## Main Text

### Introduction

Single-cell RNA sequencing (scRNA-seq) technologies have been widely used in many different areas to study the cellular states, transcriptomic dynamic changes and intercellular communications in healthy or diseased tissues (1-3). One of the fundamental problems in scRNA-seq analysis is to identify cell-types and their associated marker genes. In a standard protocol, cell-types are identified by clustering single-cells based on a set of pre-selected genes and marker genes are detected by differentially expressed gene (DEG) analysis between the clusters. Although widely used, this protocol has two important issues to be resolved.

The first issue involves the selection of genes for clustering analysis. Current scRNA-seq technologies can profile ∼10,000 or more genes for thousands of single cells in one experiment. However, not all profiled genes contain information about cell-types. Many genes have the same or very similar expressions in different cell-types. Inclusion of these uninformative genes in the clustering can seriously influence the clustering accuracy and unnecessarily increase the computational burden. Currently, genes used for clustering are often chosen as the highly variable genes (HVG) by methods such as HVGvst (4). However, HVGs are not always clustering-informative (Fig. S1A) and, vice versa, clustering-informative genes may not be HVGs (Fig. S1B). Important cell-types thus can be difficult to be distinguished by the HVGs. Other gene selection methods (5-7) have been developed. However, similar to HVG, instead of directly selecting clustering-informative genes, they all use surrogate metrics and thus have the similar problem as HVG.

The second issue is the so-called “double-dipping” (8, 9) or “selection bias” (10) problem. The expression data are first used for clustering and then reuse for DEG detection to find marker genes of the obtained clusters using available methods such as DESeq2 (11) and EdgeR (12). The uncertainty in the clustering is often ignored in the DEG detection. However, single-cells can be misclustered, especially for cell-types with similar expression patterns. The noisy cluster labels can lead to decreased precision and increased false discoveries in DEG detection. Recently, a few novel tools (10, 13) have been developed to address this problem. However, partly because these methods are often *ad hoc* and lack statistical theories for their statistical significance assessment, these methods still tend to report many false discoveries.

To address these two computational issues, we propose Festem (<u>Fe</u>ature <u>s</u>election by <u>EM</u> test) that can accurately select clustering-informative genes before the clustering analysis and identify DEGs between clusters. We observe that the gene selection and DEG detection problems are closely related problems. Ideally, a gene selection procedure should select all DEGs for downstream clustering analyses, because DEGs are cluster-informative and non-DEGs are not, and selecting DEGs should give the most accurate clustering and cell-type identification. DEGs follow different distributions in different cell-types (heterogeneous distribution) and are clustering-informative. Non-DEGs follow the same distribution among different cell-types (homogenous distribution) and are clustering irrelevant. Festem selects the clustering-informative DEGs by performing a statistical test to determine if a gene is homogenously distributed or heterogeneously distributed. Mathematical analysis shows that the homogeneity test asymptotically follows a distribution that can be bounded by a chi-square distribution (14), which thus allows Festem effectively control the false discovery rate (FDR) for DEG detections. Extensive simulation and real data analyses reveal that clustering can be significantly improved based on Festem-selected genes, and Festem can sensitively detect DEGs with high precision. Finally, in an application to a large scRNA-seq data (122,329 cells) of intrahepatic cholangiocarcinoma (iCCA), Festem identifies two types CD8+ T cells and its associated marker genes that may serve as important prognostic biomarkers of iCCA patients.

## Results

### The Festem model and algorithm

For each gene, Festem performs a statistical test to determine if its expression is homogenously distributed (not a DEG) or heterogeneously distributed (a DEG) and assigns a p-value based on the chi-square distribution 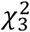 (Fig. 1). The p-values are adjusted for multiple comparisons using the Benjamini-Hochberg Procedure (15). The genes are ordered by the adjusted p-values and top genes are selected for clustering analysis. Users can then use these genes to cluster the single cells with available methods such as kmeans (16) or Louvain (17). The genes with significant adjusted p-values are reported as DEGs. Given cluster labels, Festem assigns DEGs to the clusters as their markers by the Scott-Knott test (18) (Fig. 1).

**Fig. 1.**
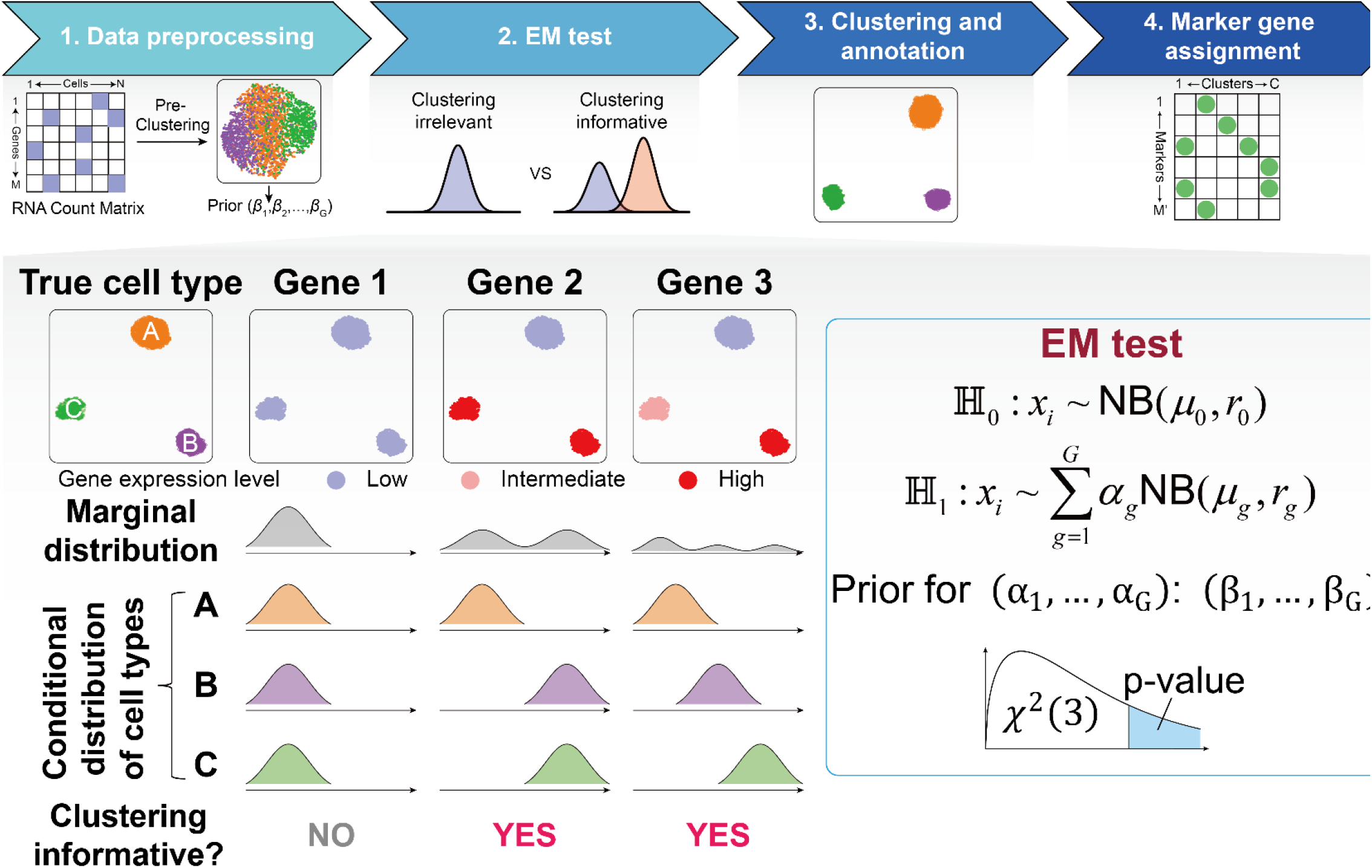
Overview of Festem. Single-cells are first roughly clustered to obtain a prior for the cell type compositions (optional). Then, Festem performs the EM-test to determine if genes are clustering informative and assigns P-values to genes based on the chi-square distribution 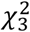 (bottom panel). Priors can be incorporated in the EM-test. Genes are ordered by the adjusted p-values and top genes are selected as DEGs and used for clustering analysis. After clustering, Festem assigns DEGs to clusters as their markers by the Scott-Knott test.

Since the cell-type labels are unknown, we cannot directly compare the distributions of gene expression in different cell-types to test whether a gene is homogenously distributed. However, if a gene is homogenously distributed, its expression distributions in different cell types are the same and identical to its marginal distribution. On the other hand, if a gene is heterogeneously distributed, it will have different distributions in at least two cell-types and its marginal distribution will be a mixture distribution (Fig. 1). Therefore, homogeneity can be tested by testing whether the distribution is a mixture distribution.

We model the expression of a gene in a cell-type following a negative binomial distribution (19). Thus, if a gene is homogenously distributed, its marginal distribution is also a negative binomial distribution; If a gene is heterogeneously distributed, its marginal distribution is a mixture of negative binomial distributions (Fig. 1). Festem tests the homogeneity using a novel Expectation-Maximization test (EM-test) for negative binomial distributions (Methods), which can be viewed as a generalized version of the EM-test for normal distributions (20). The EM-test is very similar to log likelihood ratio tests, but it avoids the pathogenicity of log likelihood ratio tests for the mixture models (20). We develop a computational efficient algorithm to calculate the EM-test statistic (Methods). It can be proved mathematically that the EM-test statistic asymptotically follows a distribution that stochastically less than 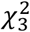 (Methods). Thus, the p-value obtained from 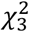 is conservative (i.e. the type I error can be effectively controlled), which is also validated by our simulation and real data analyses. Festem can also incorporate prior information, which is useful for down-weighting heterogeneously distributed genes due to factors other than cell-types such as cell cycles (21). The prior information can be very rough and may be chosen as the clustering based on the HVGs.

### Festem enables accurate clustering and DEG detection in simulation

We used simulation to evaluate the performance of Festem. In the simulation, we considered 4 different scenarios, including 2 different numbers of cell types (2 and 5 types) in the data and 2 different noise levels (low and high). For each scenario, we generated 20 simulation data, each of which had 3000 cells and 20,000 genes (Methods). We also varied the tuning parameter *G* (the number of components in the mixture distribution) to study its influence on the performance of Festem.

We first compared the clustering accuracies based on genes selected by Festem and other widely used methods including HVGvst (4), HVGdisp (22), DUBStepR (23), devianceFS (7), and TrendVar (6). For all methods, we first selected their top 500 genes and computed the first 10 principle components, then used the Louvain (17) algorithm to cluster the single cells. The adjusted Rand Index (ARI) and the Silhouette index (SI) were calculated to evaluate the clustering results given by different algorithms. Festem performed very well compared with other methods (Fig. 2A-D, Fig. S2A-D), especially in the 5-cell-type and high-noise case (Fig. 2A-D). The reason was that the available methods used surrogate metrics such as variances or deviances to select cluster informative genes. When the noise in the data is high, many selected genes just had high variances or deviances and did not contain clustering information. In comparison, Festem directly evaluated the clustering information contained in the genes and the clustering relevant genes were always more likely to be selected, regardless of the noise level.

**Fig. 2.**
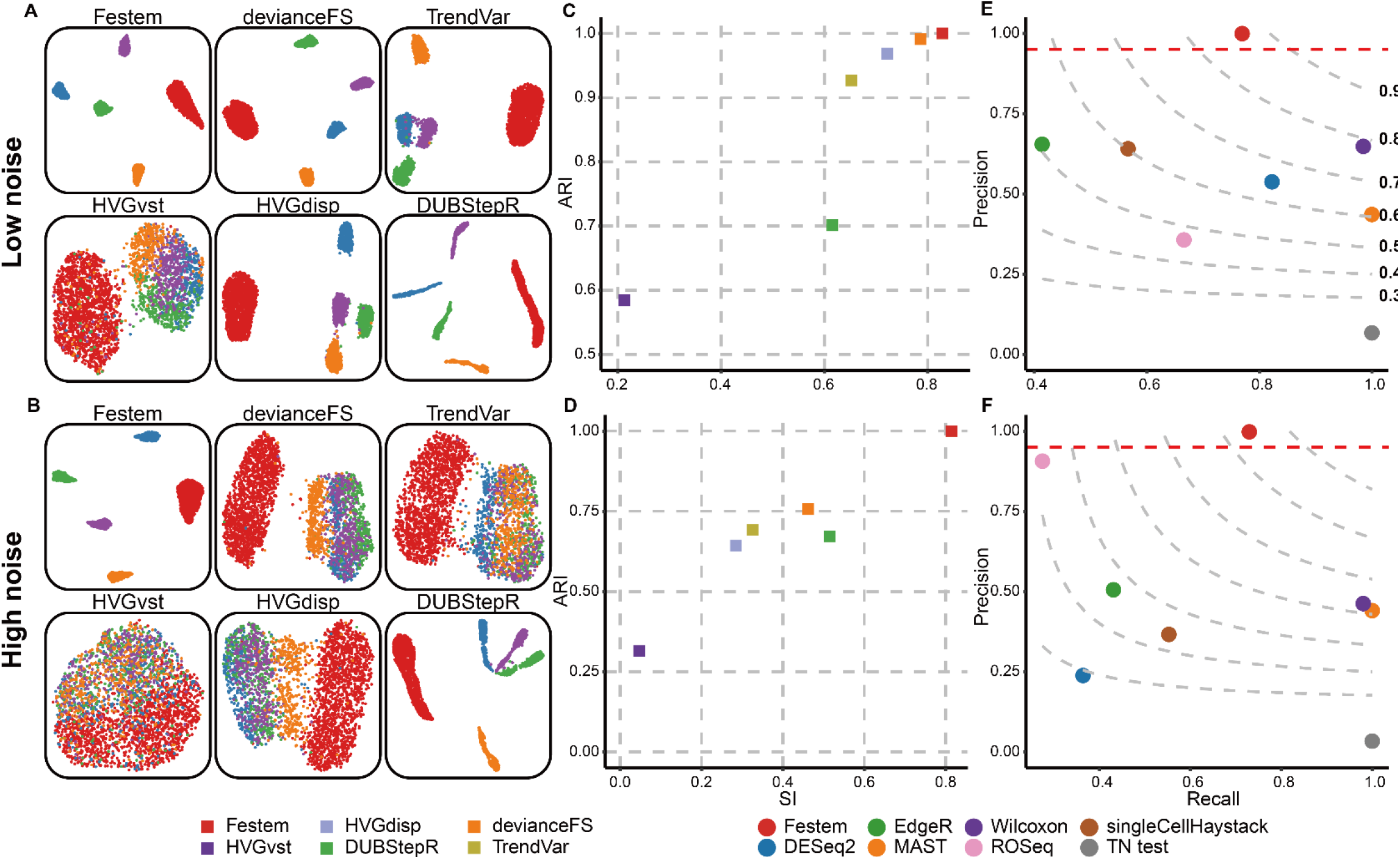
Performance of Festem on clustering and DEG analyses in simulation. Top and bottom panels correspond to simulations with low noise (400 DEGs) and high noise (200 DEGs), respectively. (**A, B**) Typical UMAP plots based on genes selected by feature selection method. Colors represent different cell types. (**C, D**) ARIs and SIs of the clustering results based on genes selected by different feature selection methods. (**E, F**) The precisions and recalls of DEG detection methods. The red lines are the nominal FDR level 0.05. The grey dashed lines are the contour lines with constant F-scores and the numbers next to the grey lines in (E) are the corresponding F-scores.

Then, we assessed the performance of Festem on DEG detection and compared with several popular and best performing DEG methods in previous benchmark studies (24, 25), including DESeq2 (11), EdgeR (12), MAST (26), the Wilcoxon rank sum test (27), ROSeq (28), singleCellHaystack (13) and TN test (10). Festem and singleCellHaystack could detect DEGs without pre-clustering of the single-cells. For other methods, we first pre-clustered the single-cells based on HVG genes and provided the clustering results to these methods for DEG detection. Festem was able to effectively control the FDR (1 - precision) and achieved reasonably good sensitivity (Fig. 2E, F and Fig. S2E, F and Fig. S3). Due to the double-dipping problem, most other methods could not control FDR and their FDRs were often much larger than the nominal level, 0.05 (the target FDR level). Although singleCellHaystack was clustering independent, it could not effectively control the FDR, possibly because of its *ad hoc* method of assessing statistical significance. ROSeq was largely able to control the FDR, but had a lower power than Festem.

Finally, we investigated the robustness and computational efficiency of Festem (Fig. S3 and S4). Overall, Festem was robust to the choice of the parameter G. Different choices of G gave very similar performances for both clustering (Fig. S4A, B, E, F) and DEG (Fig. S3A, B, E, F) analyses. In terms of computational efficiency, Festem was computationally less efficient than many gene selection methods (Fig. S4C, D, G, F), because these methods only need calculate simple statistics such as variance and deviance, while Festem need use the EM algorithm to calculate the EM-test statistic. However, comparing with the DEG methods, Festem was often computationally more efficient, especially in terms of memory usage (Fig. S3C, D, G, F). Considering that Festem could simultaneously perform gene selection and report DEGs, we concluded that Festem was comparable with other methods in computational efficiency.

### Festem had a high precision and recall in DEG detection

We evaluated the performance of Festem on DEG detection using four scRNA-seq datasets, the human peripheral blood mononuclear cells (PBMCs) data (the PBMC3K dataset, 2622 cells), the immune cells data from eight lupus patients (29) (the Kang dataset, 6410 cells) and two batches of PBMC data profiled using 3’ or 5’ 10x Genomics protocols (30) (the Zheng datasets, 8381 and 7726 cells, respectively). To compare different algorithms, we first constructed silver standards of DEGs and non-DEGs. Since housekeeping genes were usually expected to have similar expression levels in different cell types (31), we defined true non-DEGs as housekeeping genes with small Moran’s index in the 2-dimensional UMAP embedding, and true DEGs as non-housekeeping genes with large Moran’s index (Methods).

With these silver standards, we assessed the precisions and recalls of the DEG detection methods (Fig. 3A). Compared with other methods, Festem could achieve high precisions (low FDR) and high recalls, thus having high overall F-scores. Festem was also robust to the choice of the cluster number G (Fig. S5). In comparison, most other methods either had low precisions (high FDRs) or low recalls. Visual inspection showed that their reported false DEGs (housekeeping genes in the silver standard) indeed had no expression preference towards any cluster on the UMAP plot (Fig. S6). With its default settings, the popular DEG method EdgeR had high precisions but low recalls. If we turned off its *ad hoc* filters and only used corrected p-values for DEG detection, EdgeR’s recalls achieved 1 but their precisions became much smaller (Fig. S5). The other popular method DESeq2 tended to be conservative and failed due to memory limit in the two batches of the Zheng data. Because of the inaccurate clustering, the classical non-parametric Wilcoxon’s test also had low precisions. Since singleCellHaystack did not depend on clustering, singleCellHaystack performed better than other available methods. Note that the way that we built our silver standards was similar to the way that singleCellHaystack detected DEGs. Both methods relied on analyzing gene expression patterns in the 2-dimensional UMAP embedding. Thus, the evaluation might be in favor of singleCellHaystack. Methods like ROSeq, TN-test, MAST had very high recall levels, but had low precision levels.

**Fig. 3.**
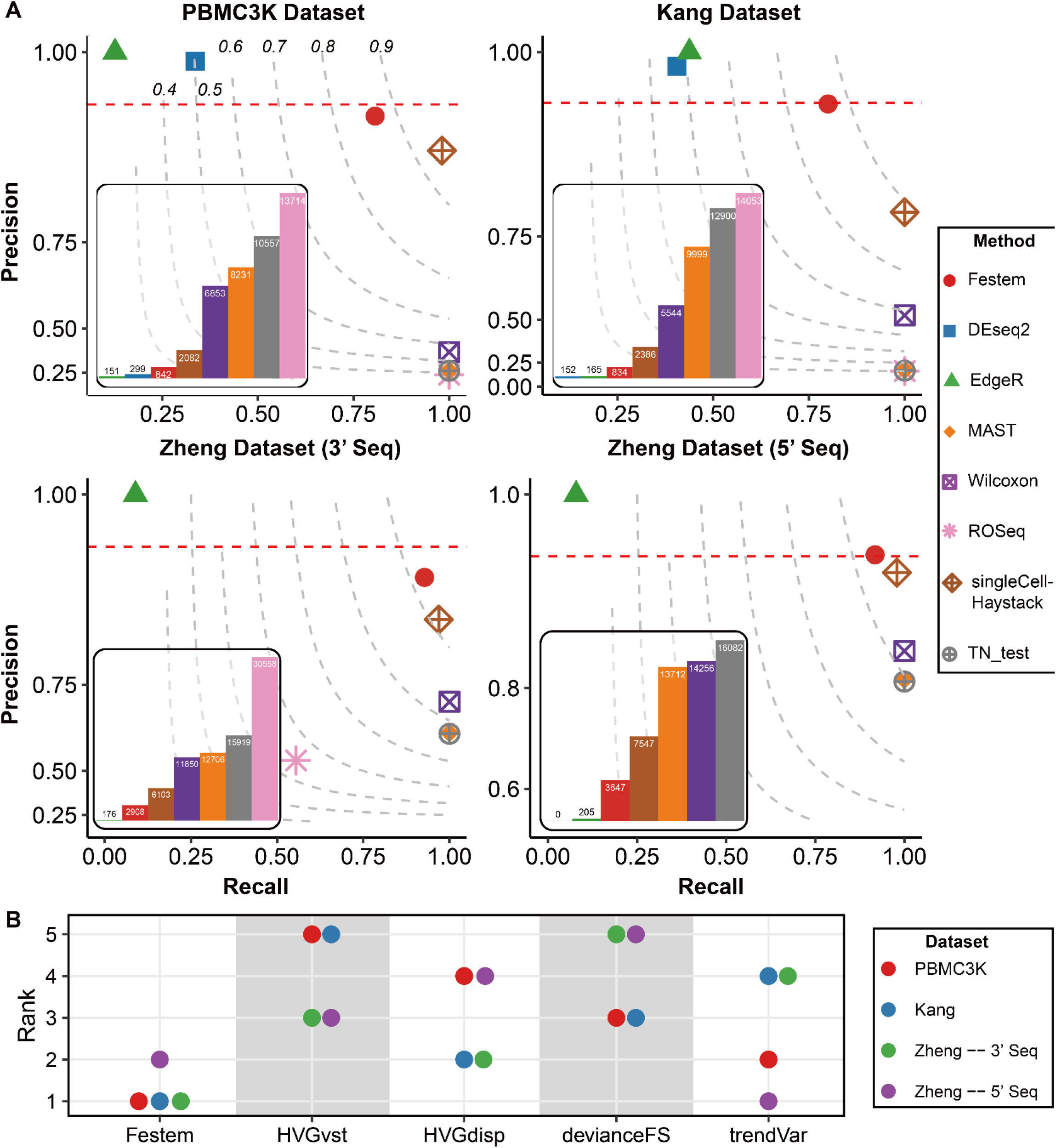
Performance of Festem on clustering and DEG analyses in real data analyses. (**A**) Precision-recall plots of DEGs reported by each method under the nominal FDR level 0.05 (red dashed lines). The grey dashed lines are the contour lines with constant F-scores (the numbers in the top-left figure). The inset bar plots are numbers of DEGs reported by each method. DESeq2 failed on the two Zheng datasets due to out of memory issue and thus were not shown. ROSeq reported no DEGs for the Zheng dataset (5’ Seq) and thus were excluded from the plot. (**B**) Ranks of feature selection methods. For each dataset, methods are ranked based on their CH indices on the corresponding UMAP coordinates.

### Festem-selected genes improve clustering

We then compared the clustering results based on Festem-selected genes with genes selected by other available methods. Due to the lack of gold standard of cell types, we employed the commonly used Calinski-Harabasz (CH) indices (32) to compare different clustering results. Larger values of the CH index corresponded to better separated clusters. Among all methods, Festem gave the largest CH index in three of the four benchmark datasets and the second largest in the other dataset (Fig. 3B), indicating Festem-selected genes could produce consistently good clustering results compared with other methods. DUBStepR was not considered in this comparison because it aimed at selecting minimum number of non-redundant genes and thus reported much less genes than other methods.

We further drew the receiver operating characteristic (ROC) curve of each method based on the silver standards constructed in the previous section and calculated the area under ROC curve (AUROC) to quantify the quality of selected genes (Fig. S7). In the four datasets, Festem always had a high AUROC (mean = 0.970), indicating that Festem stably selected most of the markers or clustering informative genes. In comparison, the AUROCs of other methods either varied greatly among different datasets or were consistently lower than that of Festem. For example, the AUROCs of HVGdisp were slightly larger than those of Festem in two of the four datasets (∼0.98), but were much smaller in the other two datasets (<0.89). Similarly, one of devianceFS’s AUROCs was slightly larger than that of Festem, but all other three AUROCs of devianceFS were much smaller. These results suggested that the superior clustering results given by Festem probably resulted from its high-quality marker gene selection.

### Festem enabled identification of often-missed fine cell types

We further investigated the cell types identified in the PBMC3K and Kang datasets based on genes selected by Festem and other available gene selection methods. We used the Louvain algorithm (17) to cluster the single cells based on the genes selected by different methods (Methods).

Using the Festem-selected genes, we identified 10 clusters in the PBMC3K data and annotated them based on the expression of canonical markers. The 10 clusters included immune cells such as naive CD4 T cells, memory CD4 T cells, CD8 T cells, CD14 monocytes, FCGR3A monocytes, Natural Killer (NK) cells, B cells and Dendritic Cells (DC) (3, 33-35) (Fig. 4A and Fig. S8). In addition to these common cell types, Festem identified a fine cell type, CD27^−^ CD4^+^ memory T cells, which were often missed by other methods (Fig. 4A).

**Fig. 4.**
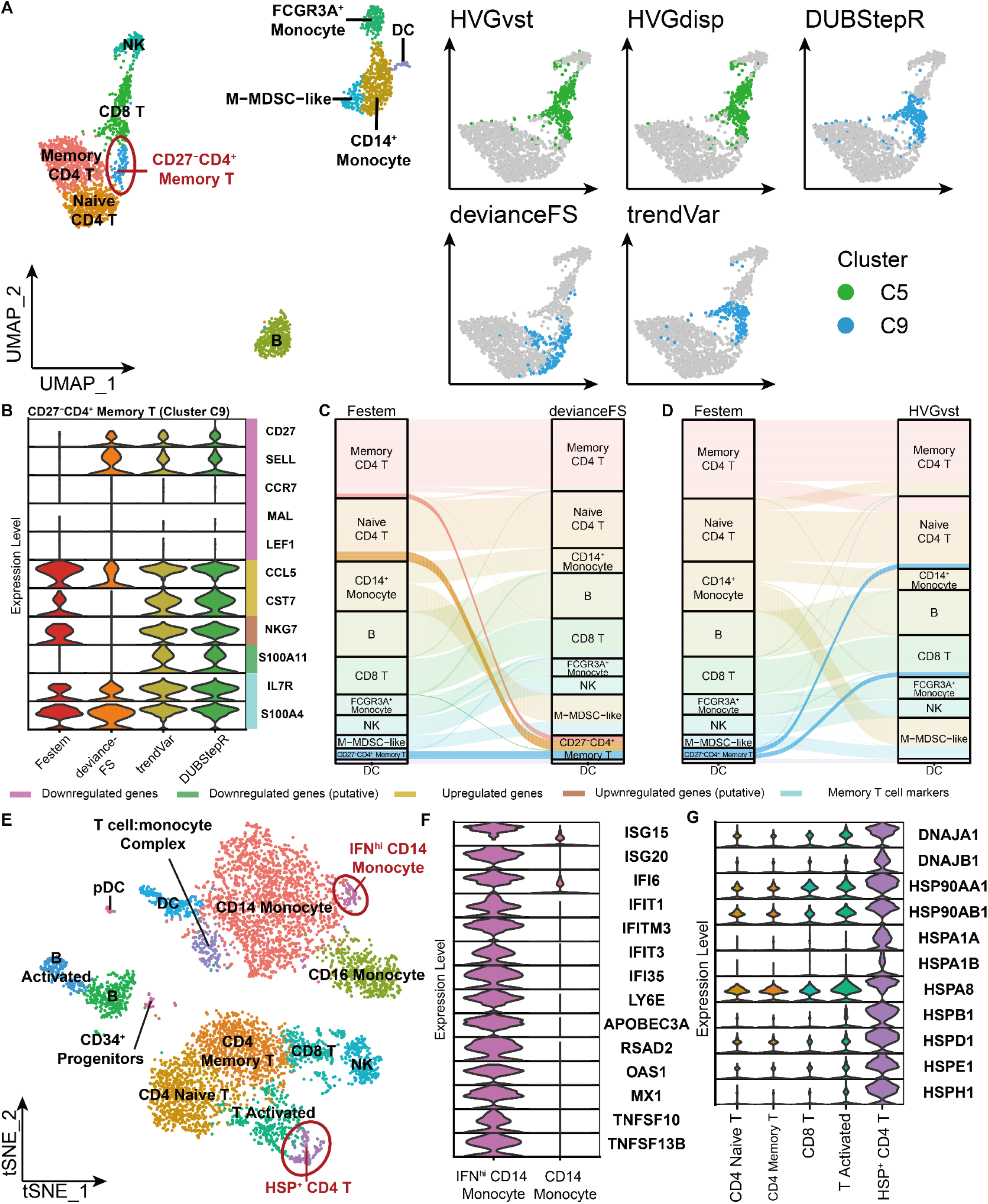
Festem identifies fine cell types in the PBMC3K and Kang dataset. (**A**) The UMAP plot of the PBMC3K dataset constructed from genes selected by Festem. Left panel: all cell types identified by Festem. Right panel: the C5 or C9 clusters by different methods (highlighted dots). These clusters are the ones identified by other methods that contain most cells of the fine cell type (CD27^−^ CD4^+^ memory T) identified by Festem. (**B**) Violin plots of CD27-CD4+ memory T cell markers. The CD27-CD4+ memory T cells identified by Festem and the C9 clusters identified by devianceFS, trendVar and DUBStepR are shown. The C5 clusters identified by HVGvst and HVGdisp are clearly mainly CD8 T cells and thus their violin plots are not shown. (**C, D**) Sankey plots for the clustering results between Festem and devianceFS or between Festem and HVGvst. (**E**) The tSNE plot of the Kang dataset constructed from genes selected by Festem. The two fine cell types often missed by other methods are highlighted in red circles. (**F**) Expressions of IFN related genes in IFN^hi^ CD14^+^ monocytes and CD14 monocytes identified by Festem. (**G**) Expressions of heat shock protein related genes in different cell types. The annotation is based on the clustering results of Festem.

The CD27^−^ CD4^+^ memory T cells identified by Festem expressed common marker genes (IL7R and S100A4) of memory T cells, but did not express *CD27*. These cells also had downregulated expression of *SELL, CCR7, MAL* and *LEF1*, and upregulated expression of *CCL5* (Fig. 4B) and thus were the CD27^−^ CD4^+^ memory T cells in the literature (36). The CD27^−^CD4^+^ memory T cells were known to be at a more differentiated state and have stronger antigen-recall responses than their CD27^+^ counterparts.

The other available methods could not correctly cluster the CD27^−^ CD4^+^ cells. For example, most of the CD27^−^ CD4^+^ cells of Festem were assigned to the C9 cluster of devianceFS (Fig. 4A, C). However, many cells in the C9 cluster of devianceFS expressed *CD27* and *SELL* or did not express *CCL5* (Fig. 4B), implying that these cells were more likely to be naïve CD4 T cells or memory CD4 T cells instead of CD27^−^ CD4^+^ memory T cells (Fig. 4C and Fig. S9). Similarly, in the clustering of HVGvst, the CD27^−^CD4^+^ memory T cells were not clustered together but were separately assigned to other cell types such as naïve CD4 T cells and CD8 T cells (Fig. 4D). One possible reason that these methods did not correctly identify the CD27^−^CD4^+^ memory T cells was that, some markers such as *CD27* and *SELL* were not selected as clustering informative genes.

For the Kang lupus dataset (29), we obtained 16 cell types based on clustering of the Festem-selected genes (Fig. 4E and Fig. S10, S11), including 2 fine cell types that were often missed by other methods. The 2 cell types were IFN^hi^ CD14^+^ monocytes with high expression of type I interferon-related (IFN) genes (Fig. 4E and F), and HSP^+^ CD4 T cells with up-regulated heat-shock-protein-related (HSP) genes (Fig. 4E and G). The HSP^+^ CD4 T cells might result from cryopreservation and thawing of T cells (37). Other than devianceFS, other available methods could not identify these HSP^+^ CD4 T cells (Fig. S11).

It was found that genes in IFN pathways were often upregulated in peripheral blood mononuclear cells of lupus and associated with the severity of lupus (38), and their upregulation was mainly in monocytes and lymphocytes (39). IFN^hi^ CD14^+^ monocytes identified by Festem were thus probably the subgroups of monocytes in lupus patients with upregulated IFN genes. The Kang dataset (the control data) had a paired IFN-stimulated single cell data. In these IFN-stimulated cells, almost all CD14+ monocytes had a high IFN expression score (Fig. S12). Joint analysis showed that, IFN^hi^ CD14^+^ monocytes from the control and IFN-stimulated groups mixed together (Fig. S13), indicating that they were the same type of monocytes that responded to type I IFN. DUBStepR and devianceFS could not distinguish the IFN^hi^ CD14^+^ monocytes and they mixed these monocytes to other types of monocytes (Fig. S11) that did not express IFN-related genes (Fig. S14).

### Festem identified new prognostic biomarkers for intrahepatic cholangiocarcinoma

We applied Festem to analyze 122,329 cells from 14 intrahepatic cholangiocarcinoma (iCCA) patients (2). We obtained 21 clusters including several subtypes of T cells, NK cells, macrophages, DC cells, B cells, fibroblasts and endothelial cells (Fig. 5A and Fig. S15). Among them, CD8 T cells had the most diverse subtypes, including terminally differentiated effector memory or effector T cells (T_emra_ cells), terminal exhausted T cells (terminal T_ex_ cells), granzyme K–positive (GZMK+) effector memory T cells (GZMK T_em_ cells) and interferon-stimulated genes (ISG)–positive T-like cells (ISG^+^ CD8 T-like cells) (40) (Fig. 5B). In addition to known markers of T_emra_ cells and terminal T_ex_ cells (Fig. 5B), Festem also identified a number of novel markers of terminal T_ex_ cells in iCCA, such as FAM3C and DUSP4. Other than devianceFS, all other available methods could not identify T_emra_ and terminal T_ex_ cells simultaneously (Fig. S16).

**Fig. 5.**
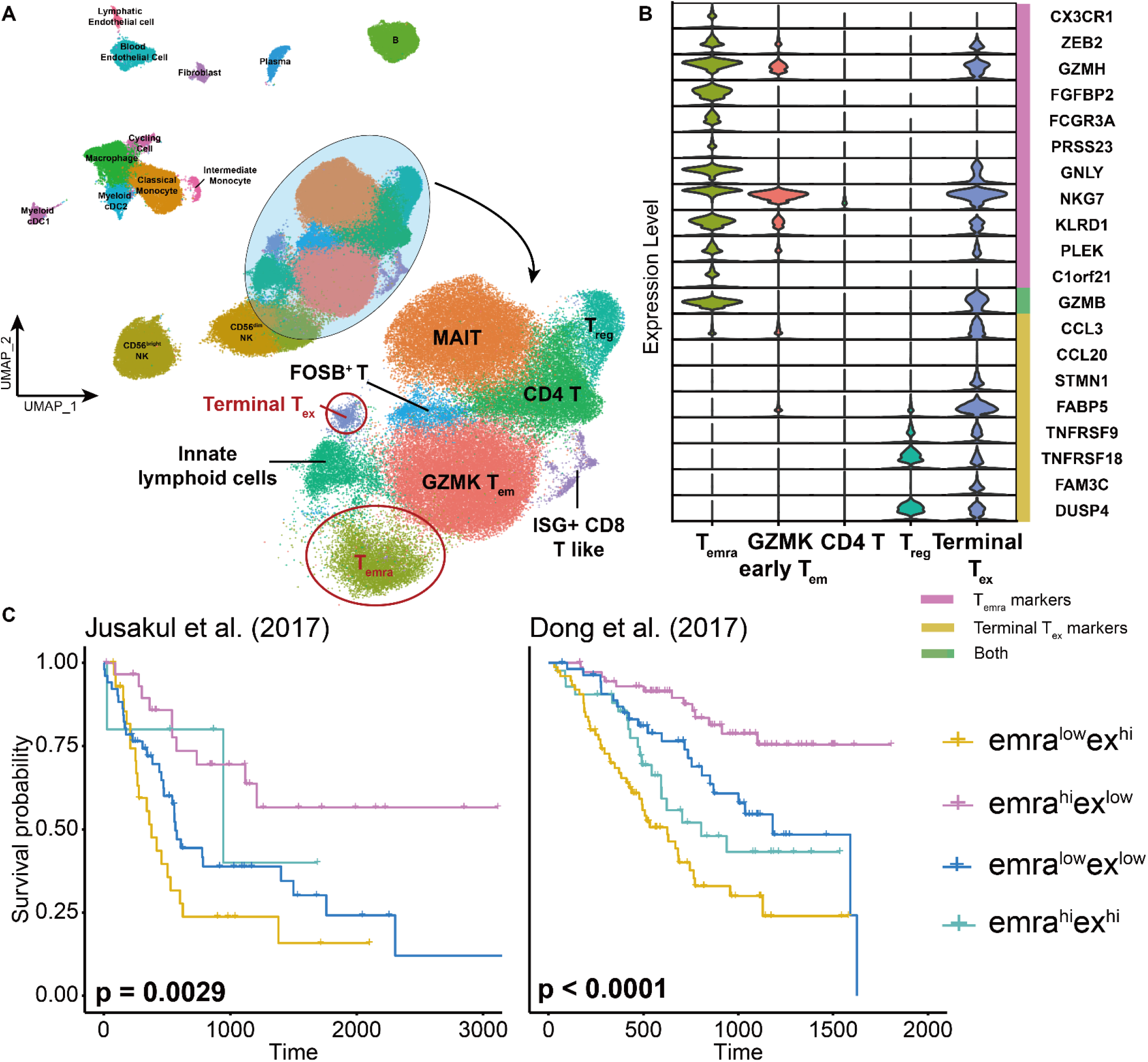
Festem identifies clinically relevant markers in the iCCA dataset. (**A**) The UMAP plot constructed from genes selected by Festem. The Festem-identified cell types are shown in different colors. (**B**) Violin plots of T_emra_ and terminal T_ex_ cell markers. (**C**) The Kaplan-Meier curves of the four groups of patients defined by the expression of markers of T_emra_ and terminal T_ex_ cells in two independent datasets. P-values given by the log-rank test are shown in the plots.

devianceFS was able to identity the two CD8 cell types, but also assigned many cells possibly of other cell types to the two CD8 cell types (Fig. S16B), leading to the decreased fold change of the marker genes of these two cell types (Fig. S16C).

We observed that the proportions of T_emra_ and terminal T_ex_ cells varied greatly among different patients (Fig. S17), and hypothesized that enrichment of T_emra_ and terminal T_ex_ cells might be predictive of patient’s survival. We therefore investigated two public iCCA datasets (41, 42) with bulk RNA-seq data and patient’s outcome data. Using markers of T_emra_ (CX3CR1) and terminal T_ex_ (FAM3C and DUSP4) cells, we grouped the patients into 4 groups, emra^high^ex^low^, emra^high^ex^high^, emra^low^ex^low^ and emra^low^ex^high^ (Methods), and found that the overall survival of patients in these groups were highly significantly different for both iCCA datasets (Fig. 5C, log rank test). As expected, the emra^high^ex^low^ group had the best survival and the emra^low^ex^high^ group had the worst survival in the 4 groups. Furthermore, using spatial transcriptomics data of hepatocellular carcinoma (43), we found that T_emra_ cells tended to be colocalized with fibroblasts in some patients (Fig. S18A). Considering that cancer-associated fibroblasts were associated with poor prognosis (44), we sought to study whether the two emra^high^ groups could be further decomposed into subtypes with distinct survival. Using the marker gene POSTN of fibroblasts identified by Festem, patients enriched with fibroblasts (POSTN^high^) indeed had significantly worse survival than other patients in both emra^high^ex^low^ and emra^high^ex^high^ iCCAs (Fig. S18B).

## Discussion

Gene selection and DEG analysis are the two most important and most commonly-used analyses in single-cell studies. The two problems are closely related and selecting DEGs should give the most accurate clustering and cell-type identification. However, because cell-types are unknown, genes are often selected using criteria that may not be directly related with clustering, and DEG analysis is performed assuming that cell-types are known. This standard protocol could miss important genes, leave important cell-type unidentified and give many false DEGs. In this paper, we present a unified framework Festem for gene selection and DEG analysis by investigating whether a gene has a homogeneous or heterogeneous distribution. Extensive simulation and real data analyses demonstrate that Fetem can improve clustering and sensitively detect DEGs with low false discoveries. Further theoretical analyses also guarantee that Festem can correctly select clustering-informative genes and control the FDR of DEG detection (14).

Festem has a tuning parameter, the number of components *G* in the mixture distribution. Theoretically, p-values given by Festem are always valid (that is, type I error can be effectively controlled) irrespective of the choice of G. Different choices of *G* only influence the power of Festem. Choosing *G* smaller than the true cluster number will lead to a decreased sensitivity. However, our empirical studies show that Festem is largely robust to the choice of *G* (Fig. S3-S5). In practice, we often can have a rough estimate of *G* and could choose *G* to be slightly larger than the rough estimate.

In the benchmarking analyses using real scRNA-seq data, the FDRs of Festem are slightly larger than the target level (Fig. 3). The reason is that Festem detected genes that are unrelated with cell-types but with other factors such as cell cycles. For example, *HMGB2, MAP1LC3B* and *NDUFA2* were non-DEGs in our silver standard (Fig. S19A), but were reported as DEGs by Festem. *HMGB2* was associated with the G2M phase (45) and its expression was significantly different in different cell cycle stages (Fig. S19B), and *MAP1LC3B* and *NDUFA2* were identified as differentially expressed between different cell cycle stages by Seurat’s function *CellCycleScoring*. Marginally, these genes were also heterogeneously distributed but the latent clusters were not cell-types but were cell cycle stages. Although we used pre-clustering priors to down-weight heterogeneously-distributed genes unrelated with cell-types, some still had highly significant p-values and were reported as DEGs. These false positives could be removed by comparing with a prior gene list that contains known genes related with cell cycles.

By analyzing a large iCCA scRNA-seq dataset, we identified two groups of important T cells, T_emra_ and terminal T_ex_ cells. The former exhibits high killing activity, while the latter has reduced cytotoxicity (46). We found that their marker genes, *CX3CR1, FAM3C* and *DUSP4*, were predictive of iCCA patients’ survival. These genes are previously reported as biomarkers in several other cancer types (47-49), further supporting their prognostic values. On the other hand, it is worth noticing that these markers may not only express in T_emra_ and terminal T_ex_ cells. For example, *CX3CR1* and *DUSP4* are reported to be expressed in some NK cells (50, 51). Hence, the prognostic values of these genes might be an aggregated effect of several immune cell types involved in cancer responses. In addition, this iCCA dataset contains over 120,000 cells. Festem was able to finish the analyses in 27 minutes with 20 cores, demonstrating that Festem scales well to very large dataset.

## Materials and Methods

### Festem Algorithm

#### The statistical model and algorithm of Festem

##### Statistical model

Let *x*_*ij*_ be the observed expression of gene *j* in cell *i* (*i* = 1, *…, N, j =* 1, *…, M*). We assume that the expression has been normalized to account for the library size by the TMM factor (52). If the gene *j* has different expression patterns in different cell types, it is clustering informative; Otherwise, it is clustering irrelevant. Given that the cell *i* belong to the cell type *g*, assume that the expression of the gene *j* follows the distribution *f*_*ij*_(*x*). Since the cell type *g* is unknown, the expression *x*_*ij*_ of the gene *j* should follow the mixture distribution 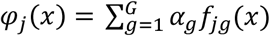, where *α*_*g*_ > 0 is the proportion of the cell type *g* in the sequenced cells 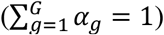. With this mixture model, if the gene *j* is clustering informative, there are at least two *g*_1_ and *g*_2_ (*g*_1_ ≠ *g*_2_) such that 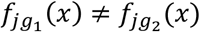. If the gene *j* is clustering irrelevant, we have *f*_*j*1_ (*x*) *= f*_*j*2_(*x*) *=* ⋯ *= f*_*j*__*G*_ (*x*) .

Studies have shown that negative binomial distribution is a suitable model for most scRNA-seq count data, especially for the mostly widely used unique molecular identifier (UMI) data (19, 53-55). Hence, we assume that *f*_*jg*_(*x*) is a negative binomial distribution. Then, the probability mass function (p.m.f.) of *x*_*ij*_ is

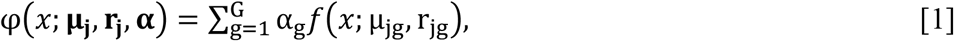

where *f*(*x*; *μ, r*) is the p.m.f. of the negative binomial distribution with mean *μ* and size *r*, 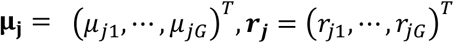 and

*α =* (*α*, ⋯ *α*)^*T*^ . Under model [1], if the gene *j* is clustering irrelevant, we have (*μ*_*j*1_, *r*_*j*1_ *=* ⋯ *=* (*μ*_*jG*_, *r*_*jG*_); Otherwise, if the gene *j* is clustering informative, we have 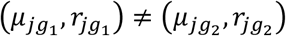 for some *g*_1_ ≠ *g*_2_. Hence, determining whether the gene is clustering informative, we can consider the following hypothesis testing problem

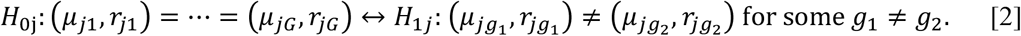

We call the null hypothesis H_0j_ the homogeneity model and the alternative hypothesis *H*_1*j*_ the heterogeneity model.

##### Testing the heterogeneity via the EM-test

We introduce the statistical test for the hypothesis testing problem [2]. Since we perform the test for each gene individually, for notation simplicity, we leave out the subscript *j* in this section. Thus, *x*_*i*_ stands for the expression of a gene in cell *i*. To quantify the evidence of heterogeneity, it is natural to use the log likelihood ratio between the alternative and the null hypothesis as the testing statistic. However, when one of *α*_*g*_ equals to zero, the Fisher’s information matrix of model (1) is not finite, leading to the failure of the log likelihood ratio test (20). To avoid this, following Li and Chen (2009) (20), we consider the following penalized likelihood:

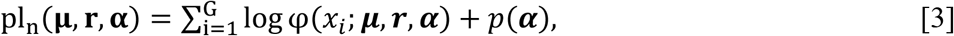

where

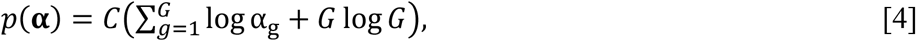

and *C* is a parameter. We always set *C* = 0.001 in all simulation and real data analyses. The penalty function *p*(***α***) is to encourage the proportions α_g_ stay away from zero.

Largely speaking, the EM-test is the difference of the maximum penalized log likelihoods [3] under the heterogeneity model *H*_1_ and the homogeneity model *H*_0_. We denote *α*_0_ *=* (1/*G*, ⋯, 1/*G*)^*T*^ and define 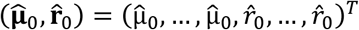 be the maximum likelihood estimation of (***μ***, ***r***) under H_0_, where

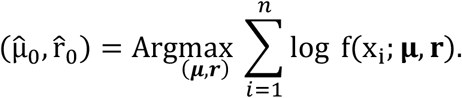

Noting that under the homogeneity model *H*_0_, the likelihood [1] does not depend on the proportion parameter **α**, then, 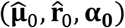 maximizes the penalized log likelihood [3] under the homogeneity model *H*_0_.

Under the heterogeneity model *H*_1_, it is difficult to directly maximize the penalized likelihood [3], and we use the expectation-maximization (EM) algorithm. We choose the initial value of the EM algorithm for (***μ***, ***r***) as 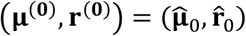. Given an initial value for the proportion parameter 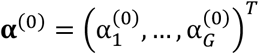, suppose that after *K* EM updates, we obtain the estimates (***μ***^(*K*)^, ***r***^(*K*)^, ***α***^(*K*)^) and define the difference of the penalized log likelihood with the initial value α_(0)_ as

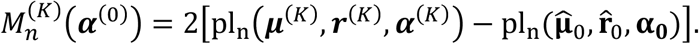

Given a set of initial values for the mixture proportions {**α**_t_: t *=* 1, *…*, T}, the EM-test statistic is defined as

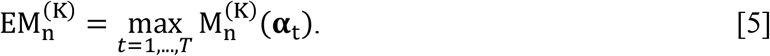

Here, we always assume that one of **α**_t_’s is **α**_0_. It can be shown that the limiting distribution of this EM-test statistic under the homogeneity model *H*_0_ is stochastically smaller than the chi-square distribution 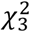. Hence, we assign p-values using the chi-square distribution 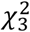. By default, Festem uses *K =* 100. We always set **α**_1_ *=* **α**_0_. Other **α**_t_’s are randomly sampled from the G-dimensional simplex. If a pre-clustering of single-cells is available, we use **α**_2_ as the proportion of different cell types in the pre-clustering.

##### The EM algorithm

We use the EM algorithm to optimize the penalized log likelihood [5]. Suppose we already have the EM estimation after k steps, **α**^(k)^ and (**μ**^(k)^, **r**^(k)^). We use the following E-step and M-step to update the parameter estimates.

- E step: calculate the weights 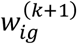(*i* = 1, …, *N*;*g* = 1, …, *G*)

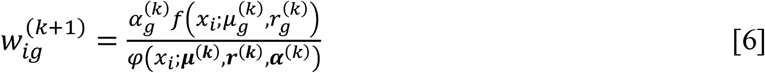
- M step:

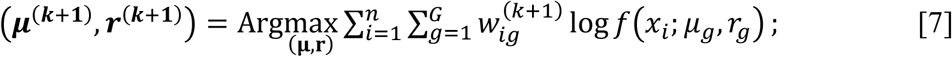

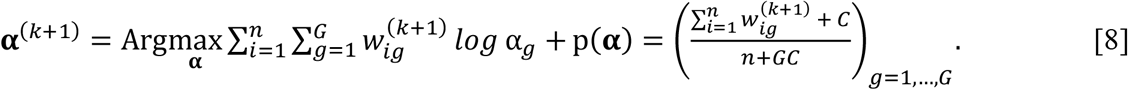

In the M-step, we optimize equation [7] with the Newton-Raphson method.

We observe that genes selected based on the EM-test statistic [5] could include genes that are related with other factors such that cell cycle or protein synthesis but unlikely to be related with cell types. Notice that 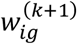 in equation [6] can be viewed as the probability of cell *i* belonging to group *g*. Therefore, to prioritize the genes related with cell types, we use a prior to reweight 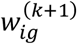. The prior is obtained by pre-clustering single-cells using HVGs. Based on the prior, we replace 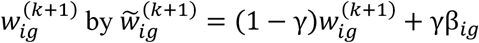, where

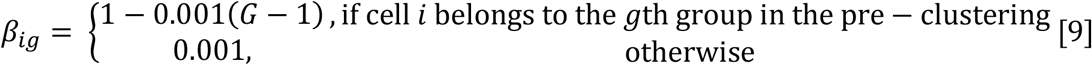

and γ is called the weight of the prior (by default = 0.05). Then, we can similarly define an EM-test statistic 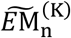. It is easy to show that 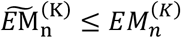 and we can still use 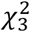 to get a valid p-value (that is, we can still effectively control the type I error).

##### DEG Reporting and Gene ranking

DEGs are reported based on the p-values and the EM statistics of the genes. More specifically, we first perform the EM-test with a very weak prior (by default *γ =* 0.05) and obtain a p-value for each gene. The prior is obtained by pre-clustering of single-cells using HVGs. Then, we adjust the p-values using the Benjamini-Hochberg Procedure. Denote the adjusted p-value for gene *j* as *p*_*j*_ . Since the prior at this step is weak, the genes with small adjusted p-values may still contain genes related with other factors but not with cell types. We further perform the EM-test with a strong prior (*γ =* 0.9). Let 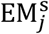 be the EM-test statistic of gene *j* with the strong prior. Given an FDR level *β*, genes with *p*_*j*_ < *β* and 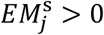 are called high confidence markers and are reported as DEGs. Genes are first ranked by the confidence of genes, and within high or low confidence gene categories, genes are ranked by the p-values *p*_*j*_ . Top genes are selected for downstream clustering and dimension reduction analyses.

##### Data preprocessing

Before applying the EM-test, we preprocess the data to remove outliers and adjust for the library size. Let *y*_*ij*_ be the observed UMI or read count of gene *j* in cell *i* (*i =* 1, *…, N, j =* 1, *…, M*). For each gene *j*, we first select cells with the largest top 5% expression *y*_*ij*_ . Let *q*_1_ and *q*_3_ be the 0.25^th^ and 0.75^th^ quantile of *y*_*ij*_ over these selected single-cells. Denote *IQR =* min{*q*_3_ − *q*_1_, 1} as the interquartile range. If any *y*_*ij*_ (*i =* 1, *…, N*) is larger than *q*_3_ *+* 3*IQR*, we consider it as an outlier. We replace the expression of the outliers with the mean of *y*_*ij*_ ‘s over the selected single-cells whose expressions are smaller than *q*_3_ *+* 3*IQR*.

After removing the outliers, we normalize the library size with the TMM factor (52). For each cell, we calculate its TMM factor by the *calcNormFactors* function in package *edgeR*. Then, we divide the count of each gene in this cell by its TMM factor and using the result as the mean of the Poisson distribution to sample a new count. Then, we apply the outlier removal procedure again to the new counts and using this count matrix to detect DEGs.

##### Assignment of DEGs to cell clusters

Note that testing hypothesis [2] does not indicate in which clusters a DEG highly expresses. After reporting the DEG list, we further use the Scott-Knott test [18] to assign each DEG to a cluster or clusters as its or their marker. For a given clustering result, which can be derived from the DEGs reported by Festem, we estimate the mean expression of a DEG in each cluster and perform Scott-Knott test with significant level 0.05 to classify these means into several expression levels, from the highest to the lowest. We call this gene the marker of a cluster if its mean is in the highest two expression levels. If we are only interested in comparing some of the clusters, we can only perform the Scott-Knott test on cells belonging to the clusters of interest.

### Comparison of feature selection and DEG detection methods on simulated datasets

#### Generation of simulated datasets

In the simulation, we generated data with 3000 cells and 20,000 genes from four different settings, with varying number of cell types and DEGs. Specifically, the four settings are: (a) 2 cell types with 400 DEGs (low-noise), (b) 2 cell types with 200 DEGs (high-noise), (c) 5 cell types with 400 DEGs (low-noise) and (d) 5 cell types with 200 DEGs (high-noise). The DEGs were set as the first *K* (200 or 400) genes. We set the DEGs only upregulated in one cell type and had the same expression distributions in other cell types.

For example, in the simulation with 2 cell types and 400 DEGs, the first 200 DEGs were set as upregulated in the first cell type and had the same distribution in the other cell type; the second 200 DEGs were set as upregulated in the second cell type and had had the same distribution in the other cell type; all other genes had the same distribution in all cells.

To make simulation data more similar to the real ones, for each gene in each cell type of the PBMC3K dataset, we first estimated the mean and size parameter of a negative binomial distribution from the count data of the gene in the cells of the cell type. The cell type annotation was chosen as the annotation of the PBMC3K dataset given in the Seurat tutorial (https://satijalab.org/seurat/articles/pbmc3k_tutorial.html#assigning-cell-type-identity-to-clusters-1). Then, we used these estimated mean and size parameters to generate simulation data. For each gene, we first randomly sampled a pair of mean *μ* and size parameters *r* from the estimated parameters. If a gene was a non-DEG, we just independently generated count data for each cell from the negative binomial distribution with the sampled parameters (*μ, r*). If a gene was a DEG, we further sampled a log_2_-foldchange *δ* from the uniform distribution in [1.5, 2.5]. For cells with the gene upregulated, we generated their expressions of the gene from the negative binomial distribution with mean *μ* · 2^*δ*^ and *r*. The expressions of the gene in other cells were generated from the negative binomial distribution with mean *μ* and *r*.

#### Benchmarking on DEG detection

We benchmarked Festem on DEG detection and compared with DEG detection algorithms DESeq2 (11), EdgeR (12), MAST (26), Wilcoxon rank-sum test (27), ROSeq (28), singleCellHaystack (13) and TN test (10). For methods requiring a clustering result, we used the following procedure to detect DEGs. In the simulation study, we first selected 1000 HVGs using HVGvst and then performed principle component analysis (PCA) dimension reduction. We then used the first 10 principle components (PCs) to construct a nearest-neighbor graph by Seurat’s *FindNeighbors* function, and clustered the cells with the Louvain algorithm (17) by the *FindClusters* function. In order to get the desired number of clusters, we applied the Louvain algorithm starting with the resolution 0.01 and gradually increased the resolution (step size 0.01). When Louvain first reported a cluster number that were the same as the true number of clusters, we stopped this process and used the corresponding cluster result. For real data analyses, we used 2000 HVGs and selected PC numbers based on the elbow plot. Clustering was performed similarly and the resolution was chosen such that the obtained cluster number was the same as the original study of the data.

For Festem, we used the prior as the pre-clustering results based on HVGs (1000 HVGs in simulations and 8000 HVGs in real analyses). singleCellHaystack does not require a clustering result but requires a dimensional reduction result. For singleCellHaystack, we first performed UMAP dimensional reduction using Seurat’s *RunUMAP* function on the 10 PC representation of the data. Then, singleCellHaystack was used to detect DEGs on both the 2D UMAP and top PC representations of the data. In simulations, we used the top 10 PCs, and in real data analyses, we selected the PC numbers based on elbow plots.

For DESeq2, we ran with default parameters and recorded adjusted p-values with and without the independent filtering step. For EdgeR, we used its quasi-likelihood test setting with default parameters. P-values both with and without the filtering step were recorded. MAST and ROSeq were also run with their default settings. The Wilcoxon rank-sum test was performed using *wilcox*.*test* function from the R language. For TN test, we first normalized the counts and split the dataset into two parts. According to TN test, one dataset was used for clustering and training of the separating hyperplanes between clusters. The clustering was performed using the same procedure described above. The other dataset was used for DEG detection based on the obtained hyperplanes.

Wilcoxon rank-sum test and ROSeq can only compare two groups of cells. When there were multiple groups, we first used these methods to do “one-versus-the-rest” comparison for each cluster and then combine the derived p-values with the Bonferroni method (56). For all methods, we adjust the derived p-values with the Benjamini-Hochberg method (15) and reported genes with adjusted p-values smaller than 0.05 as DEGs.

##### Benchmarking feature selection

We compared Festem with 5 feature selection methods HVGvst (4), HVGdisp (22), DUBStepR (23), devianceFS (7) and TrendVar (6). We used the *FindVariableFeatures* function in Seurat for HVGvst and HVGdisp. DUBStepR, devianceFS and TrendVar were run with the *DUBStepR, scry* and *scran* packages, respectively. For simulations, we selected the top 500 genes reported by each method for downstream analyses. For PBMC3K, Kang and Zheng datasets, we used the top 1000, 2500, and 3000 genes, respectively. Then, we used the first 10 PCs of the selected genes for clustering. Louvain was used for clustering and the resolution was selected as in the previous section. ARIs were calculated between the clustering results and the known cell type labels (for simulation study). UMAP dimension reduction was further performed using the 10 PC data, and SIs of known cell type labels were calculated on the UMAP coordinates by the *silhouette* function in the *cluster* package.

### Construction of silver standards of marker genes in scRNA-seq datasets

For each dataset, we selected its top 2000 HVGs, performed PCA analyses and obtained UMAP embeddings of the single-cells using the top 10 PCs. Then, we calculated each gene’s Moran index in the 2-dimensional UMAP-embedding space. The non-DEGs were taken as the housekeeping genes with absolute values of Moran’s indices smaller than a threshold *T* and with p-values larger than 0.05. For PBMC3K and Kang dataset, *T* was set to be 0.02, and for Zheng dataset, *T* was set to be 0.005. *T* was set to ensure that the expressions of genes in the non-DEG list were not visually aggregated on the UMAP plot. The DEGs were taken as non-housekeeping genes that had Moran’s indices larger than 0.1 and had non-zero expression in at least 5% cells.

## Data and code availability

All codes are publicly available at https://github.com/XiDsLab/Festem_paper. The housekeeping gene list is downloaded from http://www.housekeeping.unicamp.br/.

The PBMC3K dataset is available at https://support.10xgenomics.com/single-cell-gene-expression/datasets/1.1.0/pbmc3k/. The Zheng datasets are available at https://hub.docker.com/r/jinmiaochenlab/batch-effect-removal-benchmarking.

The Kang dataset is available in Gene Expression Omnibus (GEO) under accession number GSE96583. The iCCA dataset is available in Genome Sequence Archive in National Genomics Data Center under the accession number HRA000863. Datasets used in this study are also available at https://doi.org/10.5281/zenodo.8185811.

## Acknowledgments

This work was supported by the National Key R&D Program of China [2020YFE0204200 to R.X.], the National Natural Science Foundation of China [11471022, 71532001 to R.X.], and Sino-Russian Mathematics Center. Part of the analysis was performed on the high-performance computing platform of the Center for Life Sciences (Peking University).

## Author Contributions

R.X. conceived and supervised the study. C.W. and Z.C. formulated the method and developed the software. Z.C., C.W. and S.H. performed data analysis. S.Y. prepared and contributed to analysis of the iCCA samples. Z.C. and R.X. wrote the paper.

